# Field evaluation of non-invasive sampling for monitoring nine swine pathogens in the Republic of Korea

**DOI:** 10.1101/2025.08.29.673175

**Authors:** Hansong Chae, Serim Kim, Seung-Uk Shin, Hyunmin Shin, Gyooha Han, Mingi Han, Won Gyeong Kim, Seulki Kim, Doyoung Song, Hyuk Song

## Abstract

**Background:** Non-invasive sampling methods, such as swine oral fluid (OF) and fecal boot (FB) sampling, have gained attention due to their animal welfare considerations, ease of collection, and cost effectiveness. However, comparative data regarding non-invasive and standard sampling methods remain limited, particularly in the Republic of Korea, where research on OF and FB sampling is sparse. Therefore, we investigated representative pig pathogens, including six porcine respiratory pathogens and three gastrointestinal pathogens, using OF and FB samples for pathogen surveillance across commercial farms in the Republic of Korea.

**Methods:** The results from targeted polymerase chain reaction (PCR) assays for major respiratory and gastrointestinal pathogens obtained using OF and FB samples were compared with those obtained from standard samples. Non-invasive samples included 30 OF ropes and 30 FBs collected from nine farms, and 360 standard samples (blood, fecal, oral, and nasal swabs) were also collected. PCR assays for the nine pathogens were conducted, and the results from OF, FB, and combined OF+FB were compared with those from standard samples at the pen level.

**Results:** OF sampling showed superior test concordance with respiratory pathogens than gastrointestinal pathogens, and FB sampling demonstrated superior detection of gastrointestinal pathogens compared with OF sampling. Additionally, the combination of OF+FB sampling improved the detection concordance for all nine target pathogens investigated.

**Conclusions:** We demonstrated the feasibility and potential applicability of OF and FB sampling under field conditions in Korean pig farms. Therefore, this study supports OF and FB sampling as useful tools for swine pathogen monitoring in both research and practical field applications.

## Background

Various surveillance and diagnostic approaches have been developed to detect infectious diseases in swine. Widely used conventional sampling techniques for diagnostic tests, such as blood sampling and swabbing require individual animal restraint and can cause considerable stress to the animals [1, 2]. In contrast, non-invasive sampling methods, which do not require trained veterinarian intervention, are increasingly being adopted for improved animal welfare. These methods, including the use of cotton ropes to collect oral fluid (OF) and boot socks to collect feces (fecal boot, FB), not only simplify the process but also significantly reduce stress in the animals. Moreover, they offer significant economic and time-saving benefits, making them a prudent choice for swine farmers and veterinarians [3–6].

The OF sampling method, first described in 2008 [2, 7], leverages the natural curiosity of pigs toward novel and flexible objects. The procedure involves suspending a cotton rope to chew on in pig pens, followed by squeezing the rope to extract OF, consisting of saliva and mucosal exudate [7, 8]. In addition to the simplicity and animal friendliness of this method, as OF consists of a mixture of saliva, oral mucosal transudate, and gingival crevicular fluid, it is similar in composition to serum. Therefore, OF has been increasingly used in veterinary research to detect pathogens, such as viruses and bacteria [7]. Similarly, the FB method, introduced in 1999 [9], offers similar ease of use and non-invasiveness but primarily targets gastrointestinal pathogens, whereas the OF method is typically employed for respiratory pathogen surveillance [10]. Despite demonstrating high sensitivity in detecting gastrointestinal pathogens, particularly *Salmonella* species [5, 10], FB sampling is not as common as OF sampling.

Various diagnostic methods have been employed to examine OF and FB samples from pigs. This includes quantitative reverse transcription polymerase chain reaction (RT-qPCR), solid-phase competitive enzyme-linked immunosorbent assay (ELISA), virus or bacteria isolation, Sanger sequencing, and double antibody sandwich ELISA [11]. Several porcine pathogens, including porcine respiratory and reproductive syndrome virus (PRRSV) [12], influenza A virus in swine (IAV-S) [13], foot-and-mouth disease virus, African swine fever virus [14], porcine circovirus 2 (PCV2) [15], and *Mycoplasma hyopneumoniae* (MHP) [16] have been detected by OF samples, which are effective for diagnosis via PCR-based methods. That is, combining OF sampling with PCR-based techniques is useful for pathogen surveillance in laboratories and some field settings [3, 4].

Furthermore, compared with blood sampling or oral swabbing, which require individual restraint of animals, OF-and FB-based testing allows for simultaneous sampling from multiple animals. This method enables relatively high sampling throughput and provides valuable health data for entire swine herds. Monitoring herd-level biological data can aid in preventing the spread of pathogens by asymptomatic carriers [2]. In clinical swine practice, pooling samples has become increasingly common, with results often interpreted as representative of the age group from which the samples were collected. In this way, monitoring for pathogens through sample pooling effectively reduces veterinary diagnostic costs and facilitates testing large numbers of animals for pathogens [1, 2].

Despite the proven benefits of OF and FB sampling, research on their applicability and practicality in field conditions specific to the Republic of Korea remains limited, particularly in the context of comprehensive pathogen surveillance [6, 17–20]. Therefore, this study addresses this gap to enhance biosecurity measures in swine herds in the Republic of Korea. Pig pathogens representative of the most commonly tested pathogens in the Republic of Korea were investigated, including six porcine respiratory pathogens and three gastrointestinal pathogens [PRRSV, IAV-S, PCV2, MHP, *Actinobacillus pleuropneumoniae* (APP), *Glaesserella parasuis* (GSP), porcine epidemic diarrhea virus (PEDV), *Lawsonia intracellularis* (LI), and *Brachyspira hyodysenteriae* (BH)], using OF and FB samples across commercial Korean farms. The results were compared with those obtained through standard sampling methods (blood and nasal, oral, and fecal swabs), and a practical monitoring program was proposed for future pathogen detection on farms.

## Methods

The study was conducted in compliance with the Institutional Animal Care and Use Committee (IACUC) of Konkuk University, Republic of Korea, ensuring the ethical use of animals in research, and all swine owners entered into a written agreement with Konkuk University’s IACUC to permit ethical review of the study (under project number KU24100), and all samples were collected by trained veterinarians.

Routine blood sampling and other sample collections approved by the IACUC were performed on the animals under normal farm conditions. The animals were not euthanized as part of the study. After the completion of the study, the animals were processed according to the farm’s standard slaughter procedures, which align with local agricultural and ethical standards. No additional anesthetic or euthanasia methods were applied, as the animals followed the normal lifecycle for commercial livestock.

### Targeted pathogen selection

Targeted pathogens for this study were selected based on data from the Korea Animal Health Integrated System and the Swine Health Information Center’s Swine Disease Matrix (https://www.swinehealth.org/swine-disease-matrix/), as well as the opinions of professional veterinarians. Six porcine respiratory pathogens and three gastrointestinal pathogens were identified as significant concerns for swine health. The selected respiratory pathogens were PPRSV (PRRSV-1, PRRSV-2), IAV-S, PCV2, and MHP, which cause porcine enzootic pneumonia; APP, which causes porcine pleuropneumonia; and GSP, which causes Glässer’s disease. The selected gastrointestinal pathogens were PEDV, which causes PED; LI, which causes porcine proliferative enteropathy; and BH, which causes swine dysentery.

### Sample collection

From July 2024 to November 2024, non-invasive samples (OF and FB) and standard samples (blood, fecal swab, nasal and oral swab) were collected from nine commercial pig farms in the Republic of Korea. To minimize selection bias, three pigs were randomly selected from each pen for standard sampling regardless of clinical signs. The pigs within each pen were of similar age (growing-finishing stage, 6 to 18 weeks old) to reflect the typical herd composition in commercial swine farms (Supplementary Table 1). A total of 420 samples were obtained from nine farms in the Republic of Korea, and they were divided into 60 non-invasive samples (30 OF and 30 FB samples) and 360 standard samples (90 blood, 90 fecal swabs, 90 nasal swabs, and 90 oral swabs). The detailed collection information for each sample type are described below and in Supplementary Table 1.

To prevent cross-contamination, disposable gloves were changed between the handling of each rope and between each pen during FB sampling. Additionally, all equipment and collection materials were disposable or sterilized prior to use. All samples were transported to the laboratory on ice packs, and the average transfer time was approximately 4 h (maximum: 6 h). Upon arrival at the laboratory, samples were immediately preprocessed according to their specific characteristics and temporarily stored at 4 ℃ until DNA and RNA extraction. All extractions were completed within 24 hours of sample collection.

#### Oral fluid

A cotton rope with a thickness of 12 mm and a length of 150 cm was tied to a metal bar on the side of the pen at pig shoulder height, ensuring that the rope did not touch the floor, and left in the pen for the pigs to chew for approximately 30–40 min. OF samples were collected by placing the chewed ropes into an individual plastic zipper bag. In the laboratory, the ropes were manually squeezed into 50 mL conical tubes (SPL Life Science, the Republic of Korea). The samples were then left at 4 °C for 30 min, and 5 ml of the supernatant was carefully transferred and stored again at 4 °C until further analysis.

#### Fecal boot

FB samples were collected from pen floors using the boot-sock method (10). Two layers of disposable polypropylene shoe covers were worn over the veterinarian’s boots. During standard sampling, veterinarians walked within the pen, and the outer layer collected fecal matter and other environmental debris from the surface. The inner layer was used to avoid contamination from boots, whereas the outer layer was used for collection and testing. After sampling, the socks were placed in an individual plastic zipper bag. In the laboratory, 10 mL of PBS (Welgene Inc., the Republic of Korea) was added to the plastic zipper bag, and the contents were manually squeezed into 15 mL conical tubes (SPL Life Science, the Republic of Korea). The samples were then left at 4 °C for 30 min, and 5 ml of the supernatant was carefully transferred and stored again at 4 °C until further analysis.

#### Blood

Blood samples were randomly collected from three pigs per pen via jugular venipuncture. Pigs were appropriately restrained, and blood was drawn using a sterile needle and syringe, then transferred into tripotassium EDTA (Becton Dickinson, NJ, USA) as an anticoagulant.

Fecal, nasal, and oral swabs: Swab samples were collected from the same pigs used for blood sampling using DNA/RNA-free sterile swabs (Fisher Scientific, NH, USA). Each swab was gently inserted into the rectum, nasal cavity, or oral cavity and rotated for sample collection. In the laboratory, they were diluted in 3 mL PBS (Welgene Inc., the Republic of Korea).

### DNA and RNA extraction

Genomic DNA and total RNA were manually extracted using the AccuPrep® Genomic DNA Extraction Kit (Bioneer, the Republic of Korea) and the GeneAll® Hybrid-R™ kit (GeneAll, the Republic of Korea), respectively, according to the manufacturers’ instructions. Both DNA and RNA were extracted from 250 μL of input and eluted in 50 μL of elution buffer.

### Targeted PCR

RT-qPCR was performed for PRRSV [21], IAV-S [22], and PEDV [23]; qPCR was performed for PCV2 [24], MHP [25], LI [26], and BH [27]; and conventional PCR was performed for APP [28] and GSP [29] (Supplementary Table 2).

The RT-qPCR reaction mixture totaled 20 μL and included 10 μL of GoTaq® Probe qPCR Master Mix with dUTP (2×) (Promega, Madison, USA), 0.4 μL of RT Enzyme (50x) (Promega, Madison, USA), 1 μL of 10 μM forward primer, 1 μL of 10 μM reverse primer, 1 μL of 10 μM probe, 5 μL of RNA template, and the remaining volume of nuclease-free water. RT-qPCR was performed using the CFX Opus Real-Time PCR Systems (Bio-Rad Laboratories, CA, USA) under the following conditions: 45 ℃ for 15 min, 95 ℃ for 2 min, and 45 cycles of 95 ℃ for 15 s and 60 ℃ for 1 min. Each run included positive and negative controls.

Similarly, the qPCR reaction mixture totaled 20 μL and included 10 μL of GoTaq® Probe qPCR Master Mix with dUTP (2×), 1 μL of 10 μM forward primer, 1 μL of 10 μM reverse primer, 1 μL of 10 μM probe, 5 μL of DNA template, and the remaining volume of nuclease-free water. These reactions were also performed using the CFX Opus Real-Time PCR Systems under the same cycling conditions as the RT-qPCR, except for the omission of the 45 ℃ step.

Bio-Rad CFX Maestro 1.1 software (Bio-Rad Laboratories, Hercules, CA, USA) was used to analyze the RT-qPCR and qPCR results. Cycle threshold values were generated with a threshold of 0.1 for the target gene and 10.00% of the maximum amplitude of the sigmoidal amplification curves for positive controls.

For the PCR, the reaction mixture totaled 20 μL and included 10 μL of GoTaq® Probe qPCR Master Mix with dUTP (2×), 1 μL of 10 μM forward primer, 1 μL of 10 μM reverse primer, 5 μL of DNA template, and 3 μL of nuclease-free water. All primers were used at a concentration of 10 μM. PCRs were performed using the T100 Thermal Cycler (Bio-Rad Laboratories, CA, USA) under the following conditions: 95 ℃ for 2 min, and 45 cycles of 95 ℃ for 15 s and 60 ℃ for 1 min. After the PCR was completed, the PCR product was electrophoresed on a 2.00% agarose gel in 1× TAE buffer (Biosesang, Republic of Korea) and visualized by Midori-Green staining (Nippon Genetics, Japan) at 100 V for 20 min. The DNA separated on the gel was recorded using a GelDoc Go System (Bio-Rad Laboratories, Hercules, CA, USA).

### Analytical sensitivity

To determine the analytical sensitivity, limit of detection (LoD) tests were performed for all target pathogens. The sensitivity results of RT-qPCR for PRRSV, IAV-S, and PEDV; qPCR for PCV2, MHP, LI, and BH; and PCR for APP and GSP (Supplementary Table 2) were evaluated using 10-fold serially diluted synthetic plasmids (Bionics, Republic of Korea) for each target pathogen, ranging from 5×10^-1^ to 5×10^7^ copies/L as templates. For detection, the Ct value was considered, and the lowest concentration of synthetic plasmids that could be reproducibly detected was regarded as the presumed LoD.

The LoD was established at 50 copies/µL for PRRSV-1, PRRSV-2, IAV-S, PCV2, APP, GSP, LI, and BH and 5 copies/μL for MHP and PEDV. At the LoD concentrations, the Ct values were approximately 34–39. Consequently, we set the Ct value at 40 as the cut-off indicating positivity at Ct ≤ 40 and negativity at Ct > 40. This means Ct and Tm values were not detected in negative controls.

### Statistical analysis

The primary aim of this study was to evaluate the detection concordance between non-invasive (OF and FB) and standard sampling methods (blood, nasal swab, oral swab, and fecal swab) for pathogen detection in pigs at the pen level. Pen-level positivity was determined based on PCR results from standard samples collected from three randomly selected pigs per pen. A pen was classified as positive when at least one of the 12 standard samples (blood, nasal swab, oral swab, and fecal swab from three pigs) tested positive for a pathogen. Samples were regarded positive if the qPCR and RT-qPCR results showed a Ct value of less than 40 or if the target size (bp) was observed by electrophoresis.

The detection concordance between standard and non-invasive sampling methods was assessed using Cohen’s kappa coefficient (κ), which adjusts for agreement expected by chance [30]. Additionally, a positive rate was defined as the proportion of samples that tested positive for a specific pathogen out of the total number of samples of each type, regardless of pen-level matching. Test agreement was defined as the proportion of pens for which both the standard and non-invasive sampling methods yielded identical PCR results (either both positive or both negative). Sensitivity and specificity were defined respectively as the proportions of true positives and true negatives obtained by non-invasive sampling, using standard sampling as the reference.

All statistical analyses were conducted using R (version 4.2.2), with the epiR package (version 2.0.54) used to calculate the test agreement, sensitivity, specificity, and Cohen’s kappa with 95% confidence intervals (CIs).

## Results

In this study, detection concordance of non-invasive sampling methods (OF, FB, and combined OF+FB) was evaluated at the pen-level by comparison with standard sampling methods (Tables 1-4). As an initial step, we summarized the raw positivity rates of the nine pathogens across different sample types (Table 1), which served as a reference for the subsequent concordance analyses.

**Table 1.** Positive percentage (%) of the nine pathogens by sample type at the pen level*.

| **Sample** | **PRRSV** | **IAV-S** | **PCV2** | **MHP** | **APP** | **GSP** | **PEDV** | **LI** | **BH** |
| --- | --- | --- | --- | --- | --- | --- | --- | --- | --- |
| Oral fluid | 50.00 | 33.33 | 86.67 | 56.67 | 33.33 | 100.00 | 10.00 | 33.33 | 36.67 |
| Fecal boot | 40.00 | 23.33 | 36.67 | 43.33 | 16.67 | 63.33 | 23.33 | 26.67 | 36.67 |
| Blood | 56.66 | 39.66 | 63.34 | 10.00 | 0.00 | 0.00 | 23.33 | 0.00 | 16.66 |
| Fecal swab | 56.66 | 39.66 | 83.34 | 26.66 | 23.34 | 46.66 | 16.66 | 16.66 | 40.00 |
| Nasal swab | 46.67 | 40.00 | 73.33 | 56.67 | 50.00 | 93.33 | 13.33 | 30.00 | 40.00 |
| Oral swab | 30.00 | 43.33 | 76.67 | 50.00 | 23.33 | 100.00 | 13.33 | 40.00 | 50.00 |

**Table 2.** Test agreement, sensitivity, specificity and kappa value of pen-level comparison between OF and standard sampling methods.

| Pathogen | Test agreement | Sensitivity | Specificity | Kappa |
| --- | --- | --- | --- | --- |
| PRRSV | 76.67 | 63.33 | 90.00 | 0.53 |
|  | (64.56-85.56) | (45.51-78.13) | (74.38-96.54) | (0.33-0.74) |
| IAV-S | 76.67 | 62.50 | 92.86 | 0.54 |
|  | (59.17-88.21) | (38.64-81.52) | (68.53-98.73) | (0.26-0.82) |
| PCV2 | 86.67 | 92.59 | 33.33 | 0.26 |
|  | (70.32-94.69) | (76.63-97.94) | (6.15-79.23) | (-0.26-0.78) |
| MHP | 83.33 | 80.95 | 88.89 | 0.64 |
|  | (66.44-92.66) | (59.60-92.33) | (56.50-98.01) | (0.36-0.92) |
| APP | 60.00 | 44.44 | 83.33 | 0.25 |
|  | (42.32-75.41) | (24.56-66.28) | (55.20-95.30) | (-0.04-0.54) |
| GSP | 100.0 | 100.0 | —* | —* |
|  | (88.65-100.0) | (88.65-100.0) |  |  |
| PEDV | 73.33<br>(55.55-85.82) | 30.00<br>(10.78-60.32) | 95.00<br>(76.39-99.11) | 0.29<br>(-0.04-0.36) |
| LI | 73.33<br>(55.55-85.82) | 58.33<br>(31.95-80.67) | 83.33<br>(60.78-94.16) | 0.43<br>(0.10-0.76) |
| BH | 66.67<br>(48.78-80.77) | 50.00<br>(29.03-70.97) | 91.67<br>(64.61-98.51) | 0.38<br>(0.10-0.65) |

All values are presented with 95% confidence intervals.
\* “-“ indicates the absence of positive or negative result values, meaning no valid data were obtained.

**Table 3.**
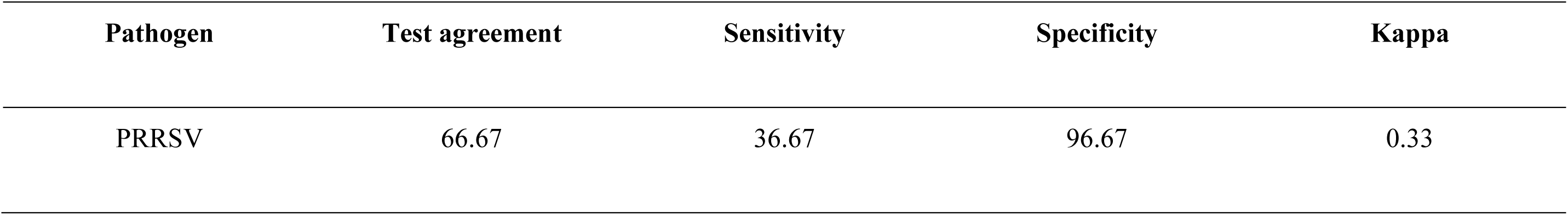
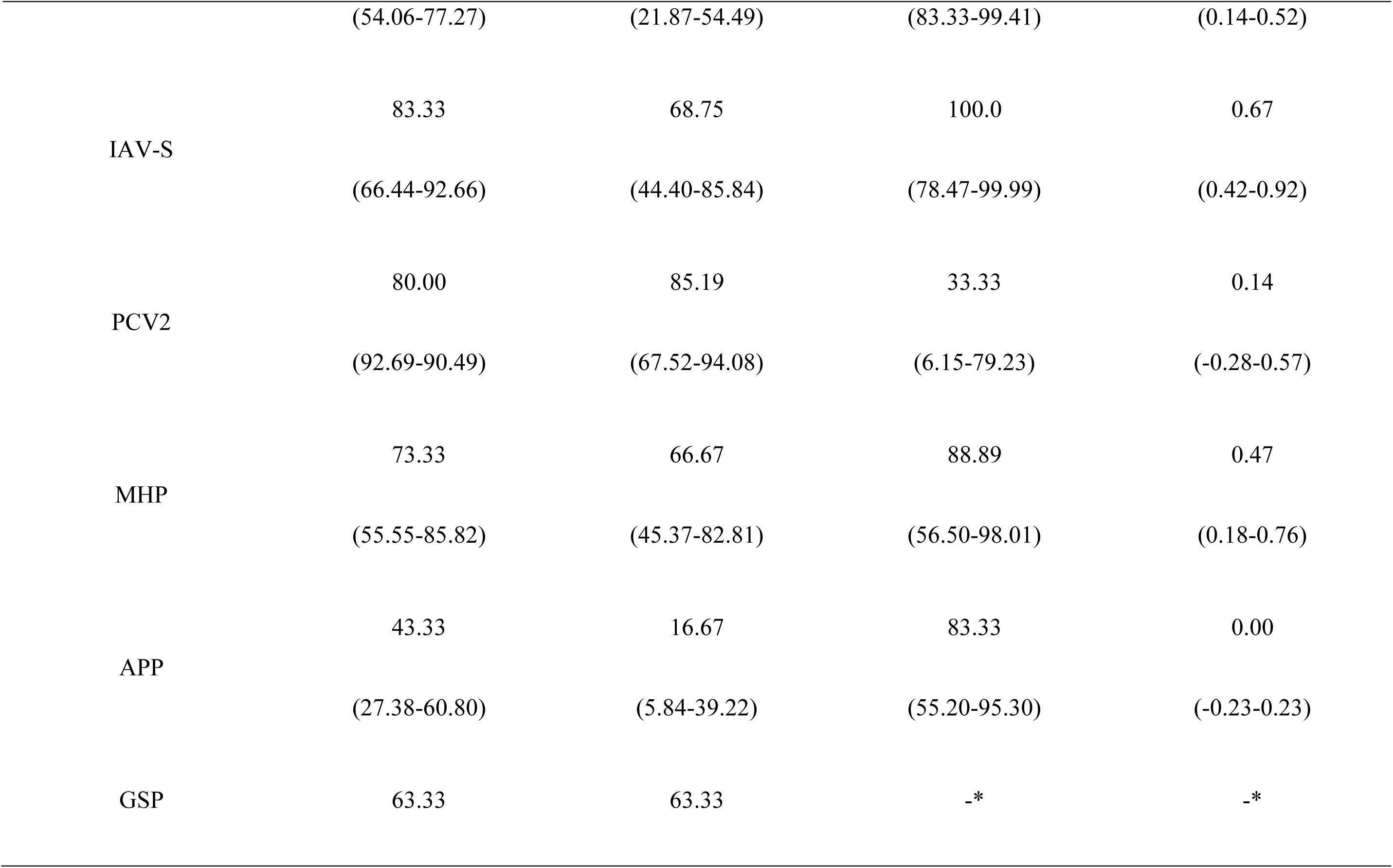
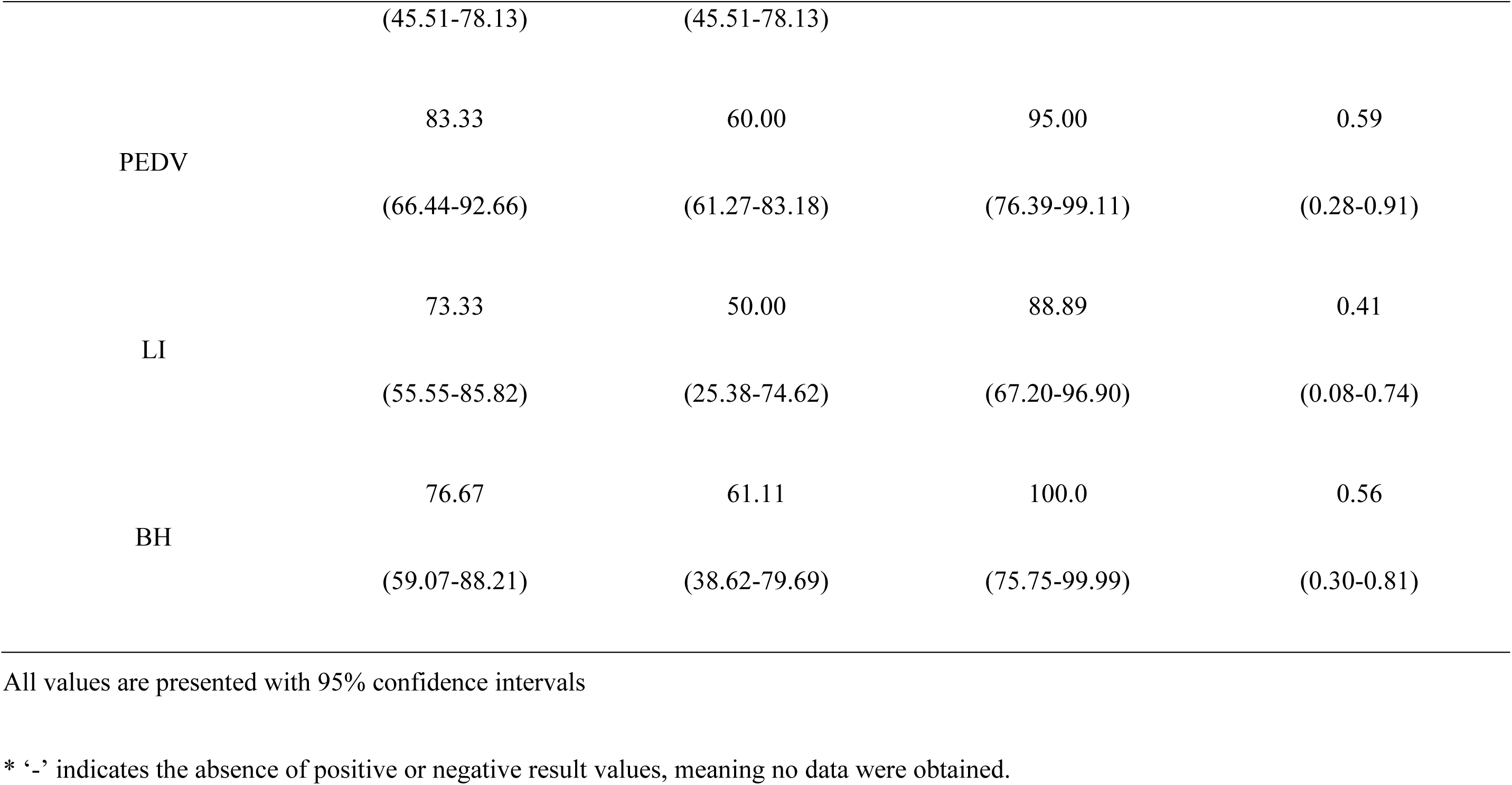
Test agreement, sensitivity, specificity and kappa value of pen-level comparison between FB and standard sampling methods.

**Table 4.** Test agreement, sensitivity, specificity and kappa value of pen-level comparison between the combined OF+FB and standard sampling methods.

| Pathogen | Test agreement | Sensitivity | Specificity | Kappa |
| --- | --- | --- | --- | --- |
| PRRSV | 78.33 | 84.00 | 74.29 | 0.57 |
|  | (66.38-86.87) | (65.34-93.60) | (57.93-85.84) | (0.36-0.77) |
| IAV-S | 86.67 | 92.86 | 81.25 | 0.73 |
|  | (70.32-94.47) | (68.53-98.73) | (57.00-93.41) | (0.49-0.97) |
| PCV2 | 90.00 | 92.85 | 50.00 | 0.35 |
|  | (74.38-96.54) | (77.35-98.02) | (9.45-90.55) | (-0.22-0.92) |
| MHP | 86.67 | 90.47 | 77.78 | 0.68 |
|  | (70.32-94.69) | (71.08-97.35) | (45.26-93.68) | (0.40-0.97) |
| APP | 63.33 | 81.82 | 52.63 | 0.30 |
|  | (45.51-78.12) | (52.30-94.86) | (31.71-72.67) | (0.01-0.60) |
| GSP | 100.0 | 100.0 | —* | —* |
|  | (88.65-100.0) | (88.65-100.0) |  |  |
| PEDV | 86.67<br>(70.32-94.69) | 87.50<br>(92.86-97.76) | 86.36<br>(66.67-95.25) | 0.68<br>(0.40-0.97) |
| LI | 73.33<br>(55.55-85.82) | 66.67<br>(39.03-86.19) | 77.78<br>(54.79-91.00) | 0.44<br>(0.12-0.77) |
| BH | 80.00<br>(62.69-90.49) | 92.86<br>(68.53-98.72) | 68.75<br>(44.40-85.58) | 0.61<br>(0.33-0.88) |

All values are presented with 95% confidence intervals
\* '-' indicates the absence of positive or negative result values, meaning no data were obtained.

Overall, the combined OF+FB sampling demonstrated improved detection concordance compared to individual OF or FB sampling and exhibited higher test agreement, sensitivity, and κ values for most pathogens. Specifically, the combined OF+FB sampling showed the highest diagnostic accuracy for detecting respiratory pathogens such as IAV-S (test agreement: 86.7%; sensitivity: 92.9%; κ = 0.73) and MHP (test agreement: 86.7%; sensitivity: 90.5%; κ = 0.68). Moreover, combined OF+FB sampling showed higher accuracy for PRRSV (test agreement: 78.3%; sensitivity: 84.0%; κ = 0.57) compared with OF sampling alone (test agreement: 76.7%; sensitivity: 63.3%; κ = 0.53).

Among gastrointestinal pathogens, PEDV and BH showed higher diagnostic sensitivity and agreement with either the FB or combined OF+FB sampling methods. Specifically, combined OF+FB sampling yielded a sensitivity of 87.5%, test agreement of 86.7%, and a κ value of 0.68 for PEDV, while for BH, sensitivity was 92.9%, agreement was 80.0%, and κ value was 0.61. FB sampling alone also demonstrated reasonable performance for gastrointestinal pathogens, notably for PEDV and BH, confirming its potential effectiveness in field applications. LI showed comparatively lower detection values, although FB or combined OF+FB sampling slightly improved detection performance compared to OF sampling alone.

High sensitivity was consistently observed for PCV2 across all sampling methods, reflecting the widespread vaccination across farms included in this study. However, specificity was markedly low (33.3%) and the κ values remained limited (κ = 0.35), indicating a reduced ability to differentiate vaccination-induced positivity from natural infections when employing these PCR-based sampling strategies. For GSP, 100% sensitivity was observed across all sampling methods, but specificity and κ values were not calculated due to the absence of negative samples, reflecting the organism’s common occurrence in swine herds as a commensal rather than as a true pathogen.

Collectively, these findings suggest that OF sampling alone provides reliable diagnostic performance for detecting respiratory pathogens while FB sampling or a combination of OF+FB is more effective for gastrointestinal pathogens under practical farm conditions. The combined OF+FB sampling strategy notably enhanced pathogen detection concordance across all categories tested, thus demonstrating its suitability for practical implementation in swine health management.

## Discussion

Blood sampling and swabbing have been the standard methods for monitoring disease levels in pig farms. However, these invasive procedures can negatively impact animal welfare and productivity by causing stress, pain, and increased risk of infection during sample collection [31]. As a result, non-invasive sampling methods such as OF and FB sampling have gained attention as practical alternatives to conventional techniques [1, 5, 6].

In this study, we evaluated the feasibility of OF and FB sampling for pathogen surveillance under commercial farm conditions in Korea by comparing pathogen detection results from standard and non-invasive samples using PCR. According to a previous study, pen-based OF sampling devices in barns housing approximately 50 pigs were able to detect PRRSV in over 80% of pens when 10% of the herd was infected [6]. Based on this evidence, we minimized our sampling to one OF sample per pen, with an average of 50 pigs per pen, aligning with previous recommendations for effective OF sampling. Sampling protocols were implemented based on established guidelines for both OF and FB methods.

Overall, combined OF+FB sampling demonstrated higher test agreement and sensitivity across most respiratory and gastrointestinal pathogens compared to either method alone, suggesting its suitability for pen-level disease surveillance. For respiratory pathogens such as PRRSV, IAV-S, PCV2, and MHP, moderate to substantial agreement was observed between OF sampling and standard sampling results. MHP showed the highest κ value (0.680) and sensitivity (86.7%) among the respiratory pathogens. PRRSV and PCV2 also exhibited acceptable diagnostic performance, with κ values of 0.570 and 0.35, and sensitivities of 70.0% and 96.0%, respectively. These findings support earlier reports suggesting that OF sampling may serve as an alternative to blood sampling for detecting PRRSV and PCV2 [6, 32, 33].

The combined use of OF and FB sampling was confirmed to be effective for the detection of various pathogens. For example, a previous study evaluating multiple individual swab types for PRRSV surveillance in weaning pigs reported sensitivities of 83.0% for ear-vein blood, 83.0% for nasal swabs, and 75.0% for oral swabs [29]. In comparison, our study demonstrated similar or higher sensitivity (approximately 84.0%) and specificity for PRRSV detection using OF samples.

While OF sampling has been well studied, FB sampling has received comparatively less attention. However, studies utilizing pen-environment swabbing and FB sampling for detecting Salmonella have shown that FB sampling can yield results comparable to OF sampling [9, 34, 35]. In our study, FB sampling showed higher test agreement and sensitivity than OF sampling alone for gastrointestinal pathogens. Specifically, PEDV and BH achieved test agreement rates of 83.3% and 76.7%, respectively, with FB samples. The combined OF+FB sampling method further improved sensitivity for these pathogens, highlighting its value for gastrointestinal pathogen monitoring.

PCV2 demonstrated high sensitivity across all sampling methods, likely due to the widespread vaccination of pigs on the participating farms. However, low specificity suggests limitations in distinguishing vaccinated animals from naturally infected ones using PCR. In addition, GSP was detected in all samples, resulting in perfect sensitivity but precluding calculation of specificity and κ values. This reflects the commensal nature of GSP rather than pathogenic presence and suggests that its consistent detection is more indicative of the appropriateness of OF sampling than of infection status.

Collectively, these findings indicate that OF sampling is particularly suitable for monitoring respiratory pathogens, while FB sampling may be more effective for gastrointestinal pathogens. Combined OF+FB sampling improved diagnostic coverage and represents an efficient and welfare-conscious approach for pen-level pathogen surveillance in swine populations.

Despite these advantages, several limitations should be acknowledged. Because pathogen concentrations may be lower in group samples such as OF and FB, the diagnostic sensitivity of these methods may be reduced for certain pathogens compared to conventional blood or swab sampling. This limitation can affect the detection of low-prevalence pathogens. Therefore, additional strategies—such as increasing the sample volume or collection frequency or combining non-invasive with conventional sampling—may be necessary to ensure comprehensive and accurate disease surveillance.

## Conclusions

This study confirmed that OF and FB sampling are practical, non-invasive alternatives to conventional sampling for swine pathogen monitoring. Among the strategies evaluated, the combined OF and FB sampling method consistently demonstrated improved diagnostic agreement and sensitivity for both respiratory and gastrointestinal pathogens. Expanding the application of these methods and optimizing sampling protocols—such as increasing sample volume or collection frequency—may contribute to the development of more sustainable and effective disease monitoring systems in the swine industry.

## Declarations

### Author’ Contributions

Conceptualization Hansong Chae; Methodology Hansong Chae; Resources Hyunmin Shin, Gyooha Han, Doyoung Song, Mingi Han and Huk Song; Validation Hansong Chae, Serim Kim, Mingi Han and Seung-Uk Shin; Data curation Hansong Chae, Serim Kim, and Won Gyeong Kim; Formal analysis Hansong Chae, Serim Kim; Investigation Seulki Kim, Seung-Uk Shin, Hyunmin Shin and Gyooha Han; Writing-original draft Hansong Chae; Writing-review & editing Hansong Chae, Serim Kim, and Huk Song. All authors read and approved of the final manuscript.

## Acknowledgements

We would like to thank Editage (www.editage.co.kr) for English language editing.

## Availability of data and materials

The original contributions presented in the study are included in the article/supplementary material, further inquiries can be directed to the corresponding author/s.

## Consent for publication

Not applicable

## Ethics approval

The study was conducted in compliance with the Institutional Animal Care and Use Committee (IACUC) of Konkuk University, the Republic of Korea, under project number KU24100, ensuring the ethical use of animals in research.

## Prior publication

A portion of this work has been presented as a conference abstract at Meeting of the Microbiology Society for Korea, but the full manuscript has not been published.

## Supplementary Tables

**S1 Table 1.** Summary of the collected pig numbers, sample types, and vaccine statuses by farm.

| Farm | No. of pens | Age of pigs* (weeks) | No. of pigs per pen | No. of group samples** |  | No. of individual samples |  |  |  | Vaccine Status*** |
| --- | --- | --- | --- | --- | --- | --- | --- | --- | --- | --- |
|  |  |  |  | OF | FB | Blood | Fecal swab | Nasal swab | Oral swab |  |
| A | 4 | 6 / 10 / 14 / 18 | 30 - 50 | 4 | 4 | 12 | 12 | 12 | 12 | PCV2 |
| B | 4 | 6 / 10 / 14 / 18 | 30 - 50 | 4 | 4 | 12 | 12 | 12 | 12 | PCV2 |
| C | 2 | 14 / 18 | 50 | 2 | 2 | 6 | 6 | 6 | 6 | PCV2 |
| D | 2 | 14 / 18 | 50 | 2 | 2 | 6 | 6 | 6 | 6 | PCV2 |
| E | 2 | 14 / 18 | 50 | 2 | 2 | 6 | 6 | 6 | 6 | PCV2 |
| F | 4 | 6 / 10 / 14/ 18 | 60 - 80 | 4 | 4 | 12 | 12 | 12 | 12 | PCV2,<br>PRRSV-2 |
| G | 4 | 6 / 10 / 14/ 18 | 60 - 80 | 4 | 4 | 12 | 12 | 12 | 12 | PCV2,<br>PRRSV-2 |
| H | 4 | 6 / 10 / 14/ 18 | 50 | 4 | 4 | 12 | 12 | 12 | 12 | PCV2 |
| I | 4 | 6 / 10 /14 / 18 | 50 | 4 | 4 | 12 | 12 | 12 | 12 | PCV2 |
| Total |  |  |  | 30 | 30 | 90 | 90 | 90 | 90 | 420 |

\*Age of pigs in each pen is shown as 'weeks/weeks/...'; each number represents a distinct pen within the same farm.
\*\*OF: oral fluid sample; FB: fecal boot sample
\*\*\* PCV2 vaccine was administered at 4 weeks of age on all farms, while PRRSV-2 vaccine was administered only on Farm F and G at 2 weeks of age

**S2 Table.**
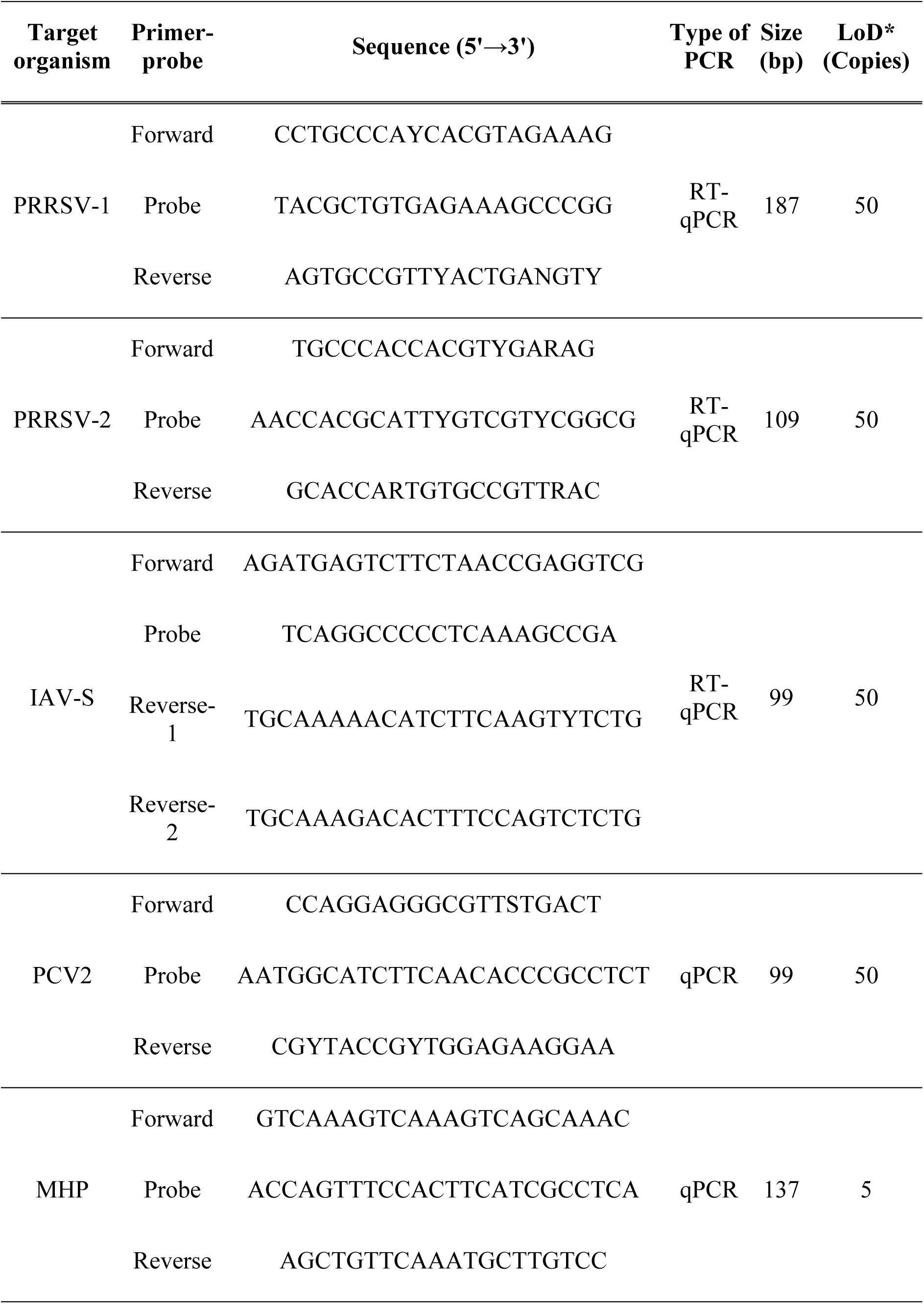
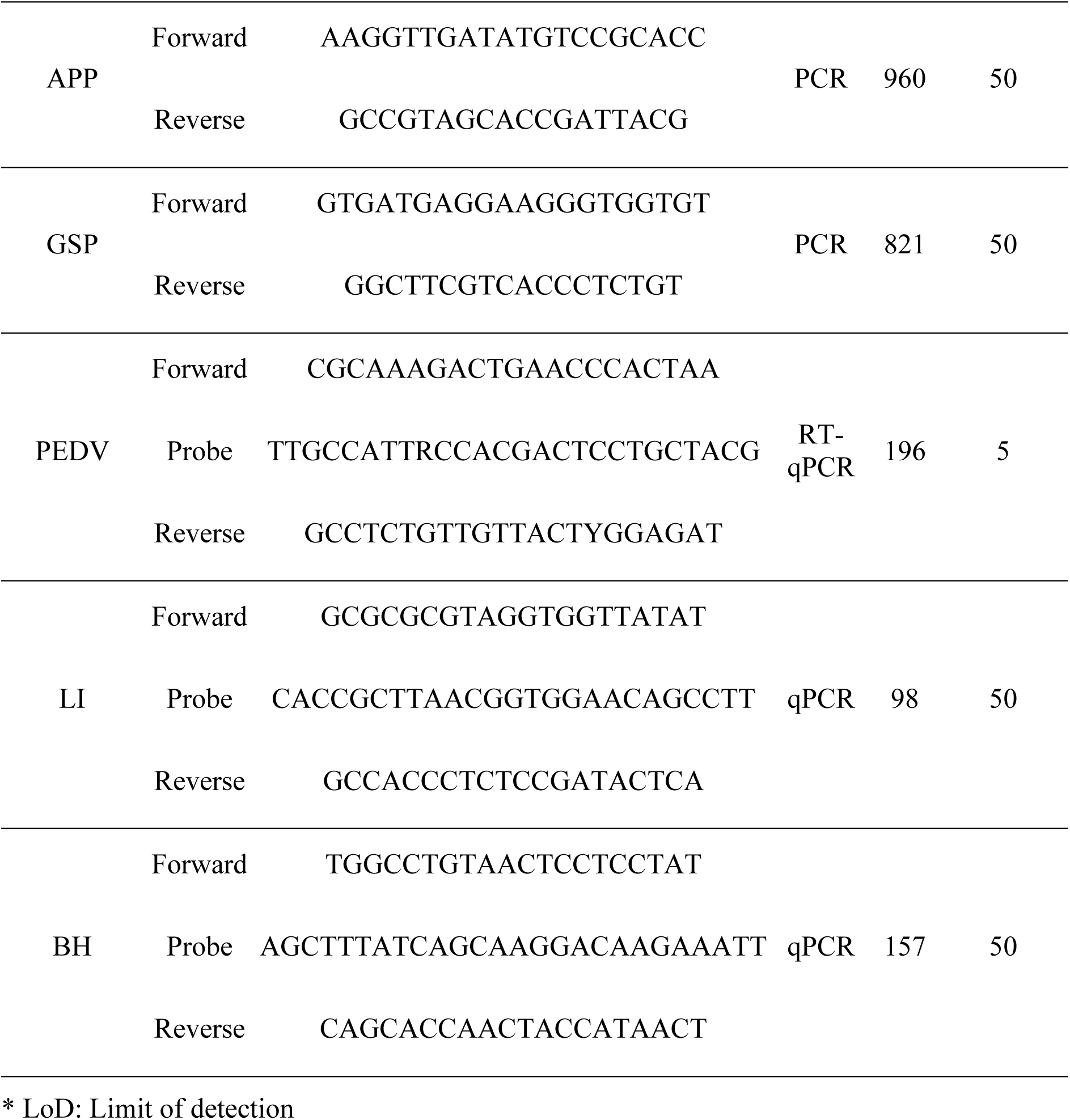
Primer-probe sets used for targeted PCRs.

